# *Chlamydia trachomatis* outbreak: when the virulence-associated genome backbone imports a prevalence-associated major antigen signature

**DOI:** 10.1101/622324

**Authors:** Vítor Borges, Dora Cordeiro, Ana Isabel Salas, Zohra Lodhia, Cristina Correia, Joana Isidro, Cândida Fernandes, Ana Maria Rodrigues, Jacinta Azevedo, João Alves, João Roxo, Miguel Rocha, Rita Côrte-Real, Luís Vieira, Maria José Borrego, João Paulo Gomes

**Affiliations:** Bioinformatics Unit, Department of Infectious Diseases, National Institute of Health, Lisbon, Portugal; National Reference Laboratory (NRL) for Curable Sexually Transmitted Infections (STIs), National Institute of Health, Lisbon, Portugal; Sexually Transmitted Diseases Clinic, Dermatovenereology Department, Centro Hospitalar e Universitário de Lisboa Central, Lisbon, Portugal; Sexually Transmitted Diseases Clinic, Lapa Health Centre, Lisbon, Portugal; CheckpointLX, Grupo de Ativistas em Tratamentos, Lisbon, Portugal; Innovation and Technology Unit, Department of Human Genetics, National Institute of Health, Lisbon, Portugal

**Keywords:** *Chlamydia trachomatis*, Lymphogranuloma venereum (LGV), recombination, *ompA*, inter-clade exchange

## Abstract

*Chlamydia trachomatis* is the most prevalent sexually transmitted bacteria worldwide and the causative agent of blinding trachoma. Strains are classified based on *ompA* genotypes, which are strongly linked with differential tissue tropism and disease outcomes. A lymphogranuloma venereum (LGV) epidemics, characterized by ulcerative proctitis, has emerged in the last two decades (mainly L2b genotype), raising high concern especially due to its circulation among men who have sex with men (MSM). Here, we report an ongoing outbreak (mostly affecting HIV-positive MSM engaging in high-risk practices) caused by an L2b strain with a rather unique genome makeup that precluded the laboratory notification of this outbreak as LGV due to its non-LGV *ompA* signature. Homologous recombination mediated the transfer of a ~4.5Kbp fragment enrolling *CT681/ompA* and neighboring genes (CT677/*rrf*, CT678/*pyrH*, CT679/*tsf*, CT680/*rpsB*) from a serovar D/Da strain likely possessing the typical T1 clade genome backbone associated with most prevalent genotypes (E and F). The hybrid L2b/D-Da strain presents the adhesin and immunodominant antigen MOMP (coded by *ompA*) with an epitope repertoire typical of non-invasive genital strains, while keeping the genome-dispersed virulence fingerprint of a classical LGV (L2b) strain. As previously reported for inter-clade *ompA* exchange among non-LGV clades, this unprecedented *C. trachomatis* genomic mosaic involving a contemporary epidemiologically and clinically relevant LGV strain may have implications on its transmission, tissue tropism and pathogenic capabilities. The emergence of such variants with epidemic and pathogenic potential highlights the need of more oriented surveillance strategies focused on capturing the C. *trachomatis* evolution in action.

## Introduction

Sexually transmitted infections (STIs) are major public health concerns globally, resulting in significant morbidity and health care costs. The World Health Organization (WHO) predicts that more than 1 million curable STIs occur each day, with an estimate of 127 million new cases of *Chlamydia trachomatis* infections in 2016 (World Health Organisation 2018). Lymphogranuloma venereum (LGV), one of the *C. trachomatis-*causing diseases of higher concern, is endemic in European countries since 2003, contrasting with the barely “inexistent” LGV cases detected until the new millennium (Nieuwenhuis et al. 2003, 2004; Savage et al. 2009; Bébéar and Barbeyrac 2009; White 2009; de Vrieze and de Vries. 2014). A real estimate of LGV infection rates in Europe is hardly traceable, not only because many LGV surveillance systems do not generate data that are considered representative of the national population (although reporting of LGV infection is compulsory in most countries with comprehensive surveillance systems), but also due to the lack of capacity of several countries to identify the LGV-causing C. *trachomatis* lineages (de Vries et al. 2019; European Centre for Disease Prevention and Control 2018). Considering that treatment of LGV differs from the recommended for ordinary urogenital chlamydial infections (Leeyaphan et al. 2016; de Vries et al. 2019), this context represents a challenge for both health care providers and public health authorities. Contrarily to the typical clinical LGV presentation (inguinal syndrome with painful lymphadenopathy), current LGV cases are essentially manifested by proctitis characterized by severe symptoms, such as anorectal pain, rectal tenesmus, haemopurulent discharge and bleeding. This “anorectal syndrome” mostly afflicts men who have sex with men (MSM), normally co-infected with HIV and other STIs (Nieuwenhuis et al. 2004; Savage et al. 2009; de Vries et al. 2019). This epidemics is being primarily caused by strains from the LGV genotype L2b (Thomson et al.2008), although some countries have been reporting increasing cases associated with the L2 genotype (Rodríguez-Domínguez et al. 2014; Peuchant et al. 2016; Isaksoon et al. 2017). Genotyping of *C. trachomatis* relies on the polymorphism of *ompA* gene, which codes for the major outer membrane protein (MOMP), its main antigen. The *ompA-* based phylogeny is incongruent with the species genome-based tree, but *ompA* genotype classification strongly correlates with tissue tropism and disease outcome, as genotypes A to C are strongly associated with the ocular trachoma, genotypes D to K with urogenital infection and genotypes L1 to L3 with LGV. This genotype grouping is generally reflected in the genome-based species tree, presenting a well-established topology with four main clades: ocular, LGV, urogenital T1 (enrolling clinically prevalent genotypes -mostly E and F) and urogenital T2 (involving less prevalent genotypes) (Harris et al. 2012; Hadfield et al. 2017). Recombination is a main driver of contemporary *C. trachomatis* evolution and diversification, mainly leading to extensive intra-clade homologous exchange of genomic regions (including *ompA*), since genetic exchange is potentiated between strains with the same tissue tropism (Gomes et al. 2007; Harris et al. 2012; Hadfield et al. 2017; Rodríguez-Domínguez et al. 2017). Inter-clade recombination is less marked. A relevant example enrolling the LGV clade concerns the report of a severe clinical case caused by an L2 strain that imported several genome-dispersed regions from a D urogenital strain, although *ompA* exchange was not observed and no further cases were reported (Sombonna et al. 2011). In fact, the inter-clade exchange of *ompA* between strains from the three disease groups (i.e., ocular, genital and LGV) is considered a rare phenomenon among contemporary clinical strains (Hadfield et al. 2017) and is believed to mediate marked tropism alterations (Andersson et al. 2016; Hadfield et al. 2017).

In the present study, we report an ongoing *C. trachomatis* outbreak caused by a recombinant strain with a rather unique genome makeup, presenting the adhesin and immunodominant antigen MOMP with an epitope repertoire typical of non-invasive genital strains, while keeping the genome-dispersed virulence fingerprint of a classical LGV (L2b) strain. This constitutes the first report of a contemporary inter-clade import of *ompA* to an LGV strain with high clinical and public health relevance.

## Methods

### *C. trachomatis* molecular diagnosis and surveillance

The National Reference Laboratory (NRL) for Curable STIs of Portugal, hosted in the National Institute of Health (NIH) Doutor Ricardo Jorge, acts as a routine laboratory providing *C. trachomatis* diagnostic to NIH clients. These are general population attending to NIH, and specific risk populations (MSM, migrants, transsexual and commercial sex workers) attending to STIs clinics. The NRL is also responsible for nationwide laboratory surveillance of *C. trachomatis.* In this context, the NRL routinely performs the molecular characterization of all C. *trachomatis* strains, either identified in the NRL itself or by any Portuguese laboratory.

*C. trachomatis* molecular diagnosis was performed by using a dual-target (chromosome *plus* plasmid) commercial nucleic acid amplification test (NAAT) (Cobas® 4800 CT/NG from Roche Diagnostics) following manufacturer’s instructions. Following the European Guidelines on the Management of LGV (de Vries et al. 2019), anorectal samples tested as *C. trachomatis* positive by the first-line NAAT test were subsequently subjected to a rapid LGV discriminatory NAAT (CLART® STIs from Genomica and/or Allplex™ Genital ulcer Assay from Seegene). Also, DNA samples from all *C. trachomatis-positive* specimens (identified either in the NRL or nationwide laboratories) are routinely subjected to *ompA* sequencing-based genotyping since LGV laboratory notification is compulsory in Portugal and requires molecular confirmation of *ompA* genotypes L1-L3 (Despacho n.^º^ 15385-A/2016; https://dre.pt/application/conteudo/105574339). Briefly, PCR and nested PCR were performed using primers NLO and NRO and primers PCTM3 and SERO2A (Lan et al. 1994), respectively, as previously described (Gomes et al. 2004), followed by nucleotide sequencing using primer *ompA-*1 (5’ TTA TGA TCG ACG GAA TTC T 3’), BigDye terminator v1.1, and capillary sequencing (3130XL Genetic Analyzer, Applied Biosystems). *ompA* genotypes are currently determined by BLASTn-based comparison (using the ABRIcate tool; https://github.com/tseemann/abricate) with a custom database enrolling reference and variant sequences of all *ompA* genotypes (Nunes et al. 2010). MEGA software (version 7; http://www.megasoftware.net) is additionally applied for fine-tuned sequence’s inspection and curation, alignment and phylogenetic reconstructions.

### Targeted *C. trachomatis* whole-genome capture and sequencing directly from clinical samples

In order to potentiate the success of the culture-independent whole-genome sequencing (WGS) of *C. trachomatis*, diagnostic DNA samples positive for the novel hybrid L2b/D-Da *ompA* genotype were preliminarily subjected to double-stranded DNA quantification (using Qubit, ThermoFisher Scientific) and subsequent absolute real-time quantification of both the number of *C. trachomatis* (targeting the single-copy *ompA*) and human genome copies (targeting β-actin), as previously described (Gomes et al. 2006). Samples selected (n = 12) for targeted WGS had a DNA amount above the minimum required for library preparation (>10ng in 7μl) and a number of *C. trachomatis* copies in the SureSelect ^XT HS^ (Agilent Technologies) input volume (7 ul) above 4 × 10^3^. Since the success of culture-independent targeted WGS does not seem to depend on the degree of human DNA content (Pinto et al. 2016), this measure did not constitute a rigid criterion (although it was taken into account to discriminate samples with low *C. trachomatis* load) (Supplemental Table S1).

*C. trachomatis* whole-genome capture and sequencing directly from selected DNA samples was performed using Agilent Technologies’ SureSelect ^XT HS^ Target Enrichment System for Illumina Paired-End Multiplexed Sequencing Library protocol (G9702-90000, version C0, September 2018, Agilent Technologies) upon enzymatic fragmentation with SureSelect ^XT HS^ and ^XT^ Low Input Enzymatic Fragmentation Kit (G9702-90050, Revision A0, September 2018, Agilent Technologies), according to manufacturer’s instructions. For this, RNA oligonucleotide baits (120 bp in size; a total of 33619 baits) were bioinformatically designed to span the *C. trachomatis* chromosome and plasmid. The custom bait design accounted for the main genetic variability among the four clades of the *C. trachomatis* species tree (Harris et al. 2012; Hadfield et al. 2017), where baits with homology to the human genome (after discontiguous megablast search against the Human Genomic + Transcript database) were excluded. After library quantification and normalization (using the Fragment Analyzer with the HS NGS Fragment kit; Advanced Analytical Technologies), *C. trachomatis* enriched libraries were subjected to cluster generation and paired-end sequencing (2 × 150 bp) in an Illumina MiSeq equipment (Illumina), according to the manufacturer’s instructions.

### Bioinformatics

In order to confirm the outbreak and characterize the genomic backbone and mosaicism of the hybrid L2b/D *C. trachomatis* strain, the following bioinformatics activities were conducted: *i*) reads’ quality analysis and cleaning/improvement, *de novo* genome assembly and post-assembly optimization using the integrative pipeline INNUca version 4.0.1 (https://github.com/B-UMMI/INNUca) (Llarena et al. 2018); *ii*) reference-based mapping and SNP/indel analysis against representative genome sequences of both the worldwide disseminated proctitis-associated *C. trachomatis* L2b strain (L2b/UCH-1/proctitis; NCBI accession numbers: AM884177.2/NC_010280.2 for chromosome and AM886279.1 for the plasmid) and the detected L2b/D *C. trachomatis* strain (strain Ct_L2b/D_PT05; ENA accession number ERZ870055) using Snippy version 4.1.0 (https://github.com/tseemann/snippy); iii) whole genome alignment and inspection using Mauve software version 2.3.1 (http://darlinglab.org/mauve/mauve.html); *iv*) coreSNP-based alignment and recombination inspection/visualization using Parsnp and Gingr tools available at Harvest suite (https://github.com/marbl/harvest). respectively (Treangen et al. 2014); v) integration of the hybrid L2b/D C. *trachomatis* strain in the *ompA*-based and genome-based species trees (Harris et al. 2012) by constructing approximately-maximum-likelihood phylogenetic trees using the double-precision mode of FastTree2 under the General Time-Reversible (GTR) model (1000 bootstraps) (Price et al. 2010); and, vi) locus-based sequence alignment and manipulation using MEGA software (version 7; http://www.megasoftware.net) (Kumar et al. 2016).

## Results

### Detection of a novel hybrid L2b/D-Da *ompA* genotype causing a potential proctitis-associated LGV outbreak

On behalf of the national laboratory surveillance of *C. trachomatis*, the Portuguese NRL for curable STIs performs *ompA* sequencing-based genotyping, not only to monitor the genetic diversity of circulating strains, but also to comply with the legislation requirements (i.e., detection of L1-L3 genotypes) for LGV laboratory notification, compulsory in Portugal since 2017. Since this year, the NRL started detecting clinical specimens tested as LGV positive by a rapid discriminatory commercial NAAT (CLART® STIs) that were not later confirmed by classic *ompA*-genotyping, as they were phylogenetically placed among D/Da genotypes. As such, we initially interpreted the incongruent results as a failure of the commercial test. However, given the continuous emergence of these discrepant cases, we retrospectively performed an alternative commercial test (Allplex™ Genital ulcer Assay) that also allows the detection of LGV strains, which confirmed the prior LGV positive results. We proceeded with a fine inspection of the *ompA*, which revealed a L2-L2b / D-Da hybrid sequence, i.e., while the first 302 bp match to *ompA* L2/L2b reference sequences (L2b/UCH-1/proctitis; GenBank accession no. AM884177.1; L2/434/BU, GenBank accession numbers AM884176.1), the region spanning the 366-1182 bp revealed a D/Da profile matching to the reference strain D/UW-3 and a Da variant (CS-908/07) circulating in Portugal (Nunes et al. 2009) (GenBank accession numbers NC_000117.1 and FJ943521, respectively). The conserved region between the 5’-3’ gene position 303-365 was estimated as the recombination crossover region.

Twenty-five cases have been detected (first case in April 2017), constituting about half (in 2017) and one third (in 2018) of the number of cases of the L2b genotype associated with worldwide proctitis-related *C. trachomatis* epidemics occurring since 2003 (data not shown). All specimens (most anorectal swabs) were collected from MSM and similar symptoms and clinical features (e.g., rectal syndrome, rectal pain, rectal tenesmus, anal discharge, rectal bleeding) were described in all cases (Figure 1). Sexually transmitted bacterial co-infections were diagnosed for most patients, especially *Neisseria gonorrhoeae* which was reported for 18 out the 25 patients. HIV status was available for 17 patients, 16 of which being positive. Multiple partners (> 10 in last 6 months) and chemsex engaging, as well as involvement with international sexual networks were also reported (Figure 1). Although all health centres where cases were detected do not have harmonized behavioural data collection procedures, hampering the identification of common sex venues or social events, known risk factors for rectal LGV, such as practices of fisting, douching and group sex in the last 12 months were reported for many MSM.

**Figure 1.**
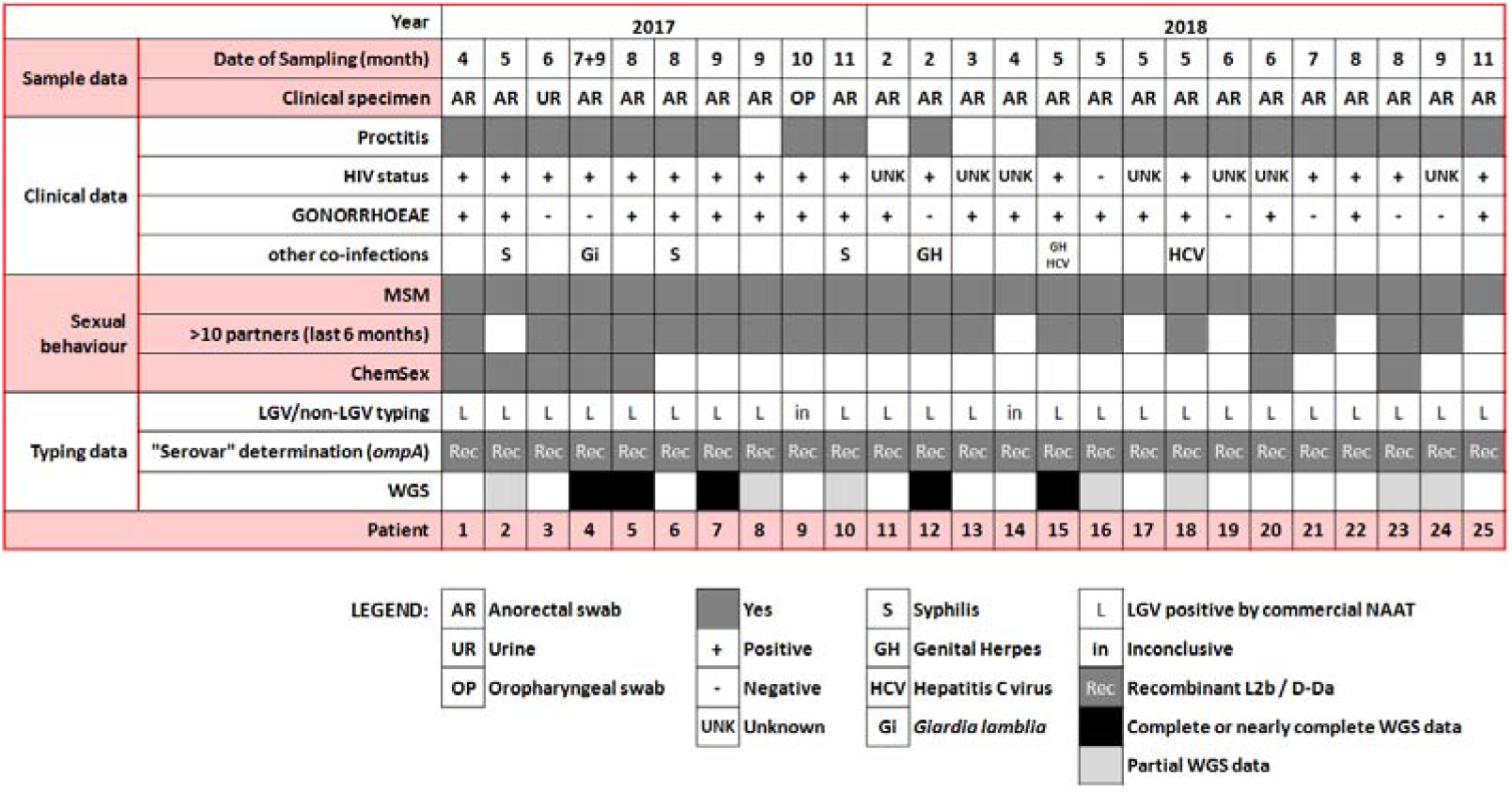
Confirmed cases of the outbreak-causing hybrid L2b/D-Da C. *trachomatis* strain. Schematic summary of the clinical data, co-infections, risk behaviors and laboratory typing results.

The real number of cases cannot be predicted considering not only the delay associated with the genotyping procedure (performed in batches and not in a real-time fashion), but also due to the frequent genotyping failures (around 30%). The last case was detected in 10^th^ November 2018.

### Genome-based outbreak confirmation

In order to confirm the outbreak and characterize the genomic backbone of the *C. trachomatis* causative strain, RNA baits-based targeted genome capture and sequencing was performed directly from diagnostic DNA samples (12 samples were selected upon bacterial load quantification through quantitative PCR). Five complete or near-complete genomes were successfully sequenced (a representative assembly was deposited in ENA under the accession number ERZ870055) (Supplemental Table S1). The phylogenetic integration of the hybrid *C. trachomatis* strain in the frame of well-established genome-based species trees (Harris et al. 2012) places it in the LGV clade, namely within the monophyletic branch reflecting the clonal expansion of the globally disseminated proctitis-associated *C. trachomatis* L2b lineage (Supplemental Figure S1A). This contrasts with its position among serovar D/Da strains in *ompA* phylogeny (Supplemental Figure S1B), and strongly supports a scenario of recombination-driven inter-clade *ompA* exchange (detailed in the next Results section) where the epidemic proctitis-associated L2b lineage was the most likely recipient strain. SNP/indel analysis of the hybrid L2b/D complete (or near-complete) genomes against the classical L2b representative genome (L2b/UCH-1/proctitis strain) confirmed the high genetic relatedness with this parental-like strain and the clonality of the hybrid strains. In fact, apart from a single high SNP density peak observed in the recombinant region (enrolling *ompA*), only 35 variant sites (involving SNPs and small indels spanned throughout the chromosome) were detected among the five hybrid L2b/D strains when compared with L2b/UCH-1/proctitis (Figure 2A; Supplemental Table S2). No differences were found in the plasmid. The L2b genomic backbone and recombination event could be also verified for the seven hybrid strains with only partial genome data by the detection of other genetic fingerprints beyond the hybrid L2-L2b/D-Da *ompA* profile detected by classical genotyping. These included the detection of SNPs/indels markers of the hybrid strain, the detection of the L2b plasmid and/or identification of sequencing reads 100% matching to the non-LGV recombinant fragment (Supplemental Tables S1 and S2). Of note, similarly to the scenario observed during the clonal expansion of the epidemic L2b lineage (Borges and Gomes. 2015), the SNP/indel-based (micro)evolutionary segregation and discreet diversification of the hybrid strain is marked by non-synonymous mutations targeting several genes potentially linked to pathoadaptation, such as genes coding for inclusion membrane proteins (e.g., CT223 and CT383), Type III secretion system effectors (e.g., CT456/Tarp) and surface proteins (e.g., CT242/*ompH, CT270/pbp3*) (Supplemental Table S2). Curiously, some clones harbor a frameshift deletion in CT157 (CTL0413 in the reference strain L2/434/Bu) gene, which codes for a Phospholipase D (PLD) protein. This represent the inactivation of an additional PLD protein, besides the other PLD pseudogenes that seem to characterize the LGV strains (Thomson et al. 2008). It is also noteworthy that several SNP/indels distinguishing the hybrid strain from the L2b/UCH-1/proctitis are homoplasic (Supplemental Table S2) as they occur in different branches of the *C. trachomatis* species tree.

**Figure 2.**
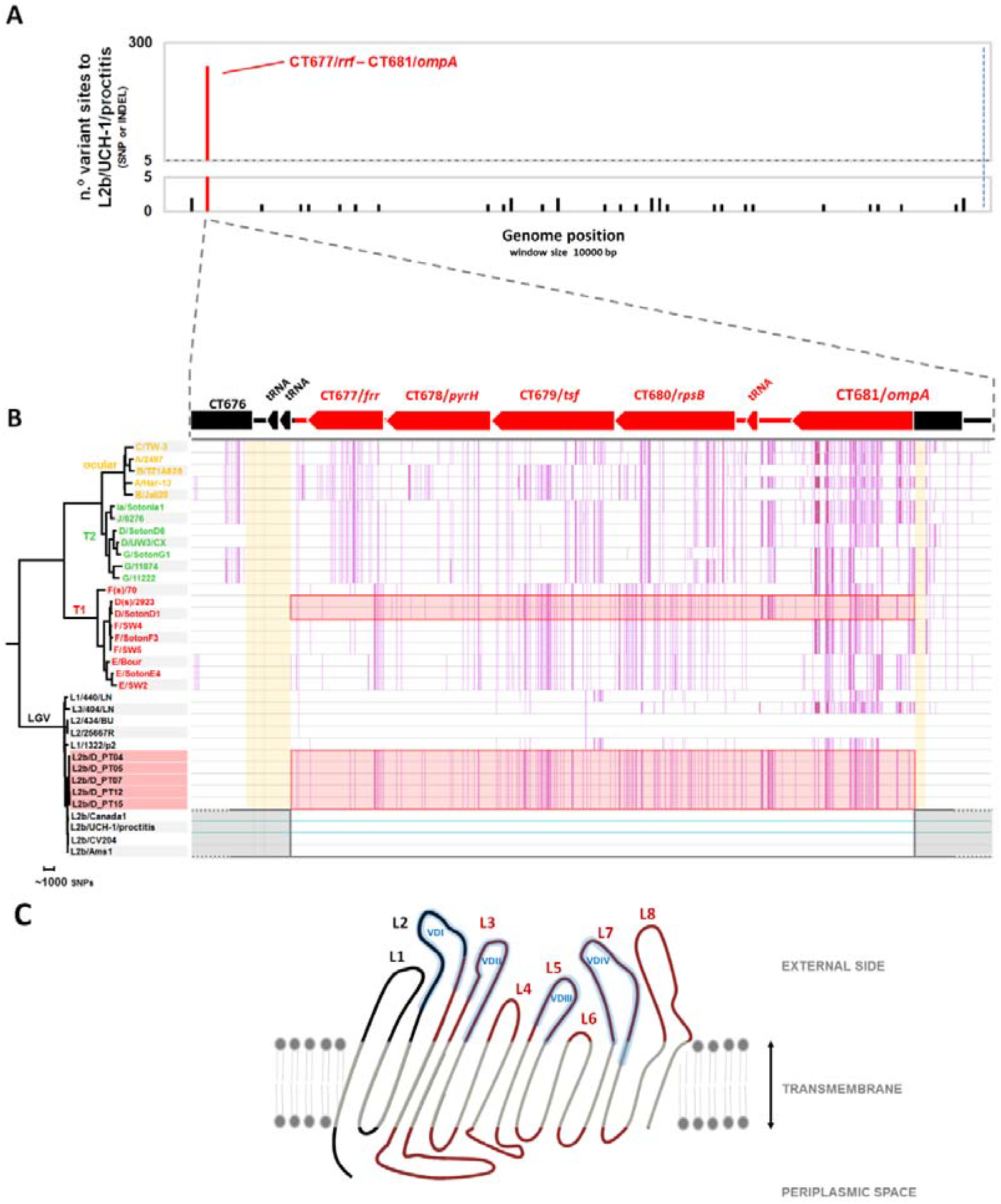
Genome backbone and mosaic structure of the outbreak-associated *C. trachomatis* L2b/D-Da recombinant strain. (A) Genome backbone and recombination analysis of the L2b/D-Da *C. trachomatis* strain. The graph shows the number of genetic differences (SNP and indels) detected among five complete or near-complete genomes of outbreak-associated L2b/D-Da clones when comparing with the genome (chromosome plus plasmid) sequence of the L2b/UCH-1/proctitis, which is a representative strain of the worldwide disseminated proctitis-associated outbreak of *C. trachomatis* L2b (a probable parental-like strain) occurring since 2003 (GenBank accession numbers: AM884177.2/NC_010280.2 for chromosome and AM886279.1 for the plasmid). Polymorphism is displayed as variant sites (SNPs or indels) (listed in Supplemental Table S2) in a window size and a step size of 10000 base pairs each, where variants introduced by the main recombination event are highlighted in red. The 100% conserved plasmid is present at the right side of the blue dashed line. **(B) Detail of the recombination event.** The L2b/D-Da *C. trachomatis* strain is a recombinant strain resulting from the genetic import of the a ~4.2Kbp genomic fragment from a non-LGV D/Da strain (most likely belonging to T1 clade) by a typical proctitis-associated L2b strain. The recombination fragment (highlighted in red by both boxes and colored genes) enrolls ^~^75% of *CJ681/ompA* and four entire genes (CT677/*frr*, CT678/*pyrH*, CT679/*tsf*, CT680/*rpsB*) (positions [55221-59461] in the L2b/UCH-1 chromosome). Estimated recombination crossovers most likely involved the two following regions (shaded in orange): i) a region with two contiguous tRNAs located between CT676 and CT677/*rrf;* and, ii) a conserved region (among serogroup B serovars: B/Ba, D/Da, E, L1 and L2) within *ompA* (5’-3’ gene position [303-365]). For schema simplification, the core-genome-based phylogenetic reconstruction enrolls sequences from *C. trachomatis* strains representing the main clades of the species tree (Harris et al. 2012) and from five outbreak-associated L2b/D-Da clones. Vertical pink lines reflect SNPs against the chromosome sequence of the parental-like L2b/UCH-1/proctitis strain (accession numbers AM884177.2/NC_010280.2) (highlighted by horizontal light blue lines). Gene orientation and annotation refers to the firstly released C. *trachomatis* genome, D/UW-3/CX strain (GenBank accession number NC_000117) **(C) Schematic representation of the topology sketch of the recombinant Major Outer Membrane Protein (MOMP)**, where the L2/L2b-like and D/Da-like non-transmembrane parts of the protein are colored in black and red, respectively. MOMP is oriented from N-terminal to C-terminal (right to left). L1-L8 indicates protein loops (L) facing the outside, and include the four variable domains (VD I-IV) of MOMP (shaded in Blue). Amino acid sequence is detailed in Supplemental Figure S2. MOMP structure was drawn based on data from Wang et al. 2005 and Nunes et al. 2010.

### Characterization of the genome mosaicism of the outbreak-related L2b/D-Da recombinant

Detailed analysis of the detected genetic mosaicism supports that the outbreak-causing L2b/D-Da hybrid *C. trachomatis* strain resulted from the genetic import of a ~4.2Kbp genomic fragment (transferring 240 SNPs, 76 of which within *ompA*) from a non-LGV D/Da strain by a typical proctitis-associated L2b strain (Figure 2A and 2B). The non-LGV parental strain was most likely a D/Da strain with E/F-like genomic backbone, i.e., belonging to clade T1 of the species tree (which involves prevalent genotypes) (Figure 2B and Supplemental Figure S1C). In fact, although NCBI BLAST analyses of the imported fragment found no single genome with 100% similarity, the “top” hits (query cover = 100%; percent identity > 99.75%) are serovar D/Da strains reported to belong to clade T1: D/13-96 (Putman et al. 2013; Hadfield et al. 2017), D/SotonD1 (Harris et al. 2012), D/SQ29 and D/SQ32 (Suchland et al. 2017). The T1 donor backbone is also strongly supported by the existence of multiple SNVs within the transferred region perfectly segregating T1 strains and also SNVs that are exclusive of serovar D/Da strains with E/F-like genome profile (Figure 2B and Supplemental Figure S1C). The recombinant fragment (position [55221-59461] in the L2b/UCH-1 chromosome) enrolls ~75% of CT681/*ompA*, which codes for the major *C. trachomatis* antigen (MOMP), and four entire genes (CT677/*rrf*, CT678/*pyrH*, CT679/*tsf*, CT680/*rpsB*) located downstream from *ompA.* Estimated recombination crossovers most likely involved the two following regions: i) a region with two contiguous tRNAs located downstream from CT677/*rrf*; and, ii) a conserved region (among serogroup B serovars: B/Ba, D/Da, E, L1 and L2) within *ompA* (5’-3’ gene position [303-365]). Of note, the latter breakpoint region has been also involved in the partial import of *ompA* from classical trachoma-causing strains to clade T2 strains that were shown to cause trachoma (Andersson et al. 2016) The recombinant L2b/D-Da MOMP displays a total of 19 amino acid (aa) replacements and 1-aa insertion when compared with the parental-like L2b MOMP, with 18 of the aa variants falling within surface-exposed epitope-enriched highly variable domains (VD) (Supplemental Figure S2). In fact, the recombination event yielded a new combination of epitopes in the *C. trachomatis* immunodominant antigen, MOMP, mixing serovar L2/L2b epitopes (VDI region) and serovar D/Da-like epitopes (from VDII to VDIV) (Figure 2C; Supplemental Figure S2).

## Discussion

WGS-based molecular typing is still not a reality for *C. trachomatis* routine surveillance, despite the need to timely characterize circulating types and variants in order to disclose transmission chains, guide therapeutics and identify emerging public health concerns. However, the recent history of *C. trachomatis* infections constitutes a hallmark example of how the dynamic pathogen evolution (rather than technological advances) demands important modifications in diagnosis, surveillance and therapeutic strategies. The first relevant example of the “evolution in action” with impact on molecular diagnosis / surveillance relies on the emergence of a *C. trachomatis* variant, detected in 2006 in Sweden, with a deletion precisely in the plasmid region targeted by widespread commercial nucleic acid amplification tests (Ripa and Nilsson 2006). Retrospective studies revealed that this variant was circulating at least since 2001, and displayed a rapid spread (reaching 64% of all diagnosis in few years in Sweden) due to diagnostic failure and consequent lack of treatment (Unemo et al. 2010; Unemo and Clarke 2011). This scenario posed a rapid re-design of molecular diagnostics that now target both the plasmid and the chromosome.

*The C. trachomatis* evolution should be potentiated by the increasing human population density and circulation, as well as by changes in sexual networks enhancing strains’ spread and mixing. According to this, the MSM-associated global emergence of the successful proctitis-related L2b clone implicated the modification of international guidelines for *C. trachomatis* molecular identification/typing, which have come to recommend tests targeting LGV-specific genome loci and differential treatment for LGV cases (Lanjouw et al. 2016; de Vries et al.2019). The unique recombinant character of the outbreak-causing L2b/D strain reported in the present study leads to an “erroneous” classification and potential subsequent treatment if only the traditional *ompA* typing is performed, since *ompA* phylogeny places this hybrid strain among serovar D/Da strains (Supplemental Figure S1). Despite commercial NAATs for rapid LGV / non-LGV differentiation were proficient in detecting these recombinant strains as LGV, they are not widely available and lack validation (Lanjouw et al. 2015; de Vries et al. 2019). Moreover, a laboratory sub-notification of LGV cases (compulsory in multiple countries [European Centre for Disease Prevention and Control, 2018]) would be yielded, as *ompA* genotyping is still required to accomplish with the Commission Implementing Decision (EU) 2018/945 of 22 June 2018 (https://eur-lex.europa.eu/legal-content/EN/TXT/?uri=uriserv:OJ.L_.2018.170.01.0001.01.ENG). which requires L1, L2 or L3 identification to fulfill the criteria for LGV confirmed case. For example, in Portugal, where LGV laboratory notification is compulsory and LGV case definition (Despacho n.º 15385-A/2016; https://dre.pt/application/conteudo/105574339) is in compliance to the EU legislation, these outbreak-associated cases cannot be formally notified as LGV. This has already triggered an ongoing process to revise the criteria for laboratory LGV notification. Altogether, this may trigger the spread of the hybrid outbreak strain, especially due to its circulation among MSM, most of them co-infected with HIV and other STIs, normally reporting a sexual behavior marked by chemsex engaging and, in some cases, involvement in international sexual networks. Considering this methodological and epidemiological context, one can hypothesize that the outbreak has already crossed boarders, although remaining unnoticed. This whole scenario not only highlights the need to revise laboratory methods for notification, but also sustains the international recommendations to target LGV-specific genome loci for detecting, monitoring and controlling LGV outbreaks. Several MLST-like systems capable of LGV identification are described (e.g., Klint et al. 2007; Pannekoek et al. 2008; Dean et al. 2009; Bom et al, 2011) but none of them was broadly adopted by the scientific community. Curiously, *in silico* extraction of MLST profiles using the three schemas currently available at pubMLST (https://pubmlst.org/) showed that the recombinant L2b/D-Da strain cannot be distinguished from the parental-like L2b/UCH-1/proctitis lineage by these MLST strategies [belonging to ST44 *(Chlamydiales* schema; Pannekoek et al. 2008), ST1 (*C. trachomatis* schema; Dean et al. 2009) and ST58 (*C. trachomatis* Uppsala schema; Klint et al. 2007)]. As such, the present outbreak reinforces that the advisable application of multi-loci typing strategies should always be coupled with *ompA* genotyping. From a clinical point of view, this specific epidemics reinforces that MSM (especially HIV-positive MSM) should be routinely screened for *C. trachomatis* infection (oral, urethral and anal), and voluntary partner notification and epidemiological LGV treatment should be considered due to possible sexual network clustering (sex abroad and chemsex). Also, second generation surveillance for LGV (e.g., centralized integration of both microbiological and demographic/behavioural data at the NRL, safeguarding the necessary ethical issues) would be advisable to track both “evolution” and “context” of *C. trachomatis* (namely LGV) epidemics. For example, in this outbreak, an harmonized behavioural inquiry among the enrolled STI clinics could have allowed more complete reports about practices of fisting, douching and group sex in the last 12 months, which are known risk factors for rectal LGV (MacDonald et al. 2014) that were reported by most patients from one of the clinics where this data is systematically questioned.

The inter-strain exchange of the whole or part of the *ompA* gene is a natural phenomenon in *C. trachomatis* (Dean et al. 1992; Brunham et al. 1994; Hayes et al. 1994, Gomes et al. 2007, Harris et al. 2012; Rodríguez-Domínguez et al. 2017; Anderson et al. 2016; Hadfield et al. 2017). However, besides the known ancient *ompA* diversification (leading to a *C. trachomatis ompA* genotype grouping incongruent with the genome-scale tree topology) (Harris et al. 2012), the present study constitutes the first report of an epidemiologically and clinically relevant hybrid strain presenting a classical LGV chromosome backbone while harboring most of the major antigen (MOMP) from a non-LGV strain. This evolutionary step is not surprising in the epidemiological and demographical context it emerged. In fact, the MSM-associated worldwide epidemics of L2b (occurring during the last two decades) potentiates the co-infection with high prevalent urogenital strains (e.g., T1 clade), which are usually associated with asymptomatic infections, favoring multiple rounds of infections and co-infections. Besides the main recombination event, the hybrid strain also revealed discreet genome-dispersed homoplasies that further corroborates a scenario of recurrent exchange of genetic material with other non-LGV strains, which is an microevolutionary mechanism occurring in parallel with the fixation of *de novo* mutations mostly affecting host-interacting proteins with potential role in pathoadaptation (Supplementary Table S2). The observed genome mosaicism yields, not only a new combination of MOMP epitopes (mixing D/Da-like and L2/L2b-like epitopes within MOMP) (Figure 2; Supplementary Figure S2), but also a novel antigenic composition at the whole proteome level. In fact, most MOMP epitopes (both T- and B-cell epitopes) belong to strains typically associated with non-invasive genital tropism, whereas the remaining antigen repertoire (e.g., Polymorphic membrane proteins - Pmps; Chlamydial protease-like activity factor - CPAF; Inclusion membrane proteins - Incs; OmcB) (Thomson et al. 2008; de la Maza et al, 2017) is typical of LGV strains causing anorectal and/or inguinal syndromes. A major impact on the its ability to avoid immunological protection can indeed be predicted as even modest alterations in the immunodominant MOMP may substantially change *C. trachomatis* antigenic presentation (Suchland et al. 2003; Kari et al, 2008; 2009). Besides the potential impact of this novel L2b/D-Da mosaicism on the interaction with the host immune system, other bacterial functions that may influence pathogenesis could also have been affected. In fact, the immunodominant MOMP, which alone constitutes about 60% of the membrane dry weight (Caldwell, 1981), is also a porin (Bavoil et al, 1984) and adhesin (Su et al. 1990) that is assumed as a major host-interacting bacterial factor underlying differential tissue tropism. In strong support of this, a study of *C. trachomatis* infections in Australian Aboriginal patients revealed that typical genital strains (clades T1 and T2) became capable of causing trachoma after importing MOMP (alone or together with other adhesin/antigens coded by *pmpEFGH*) from ocular strains (Andersson et al. 2016). We hypothesize that, although the selective advantage of the recombinant strain most likely relies on the novel signature of the hybrid MOMP, the other exchanged genes neighboring *ompA* (CT677/*rrf*, CT678/*pyrH*, CT679/*tsf*, CT680/*rpsB*), which play key functional roles (ribosome recycling, pyrimidine metabolism, translation elongation and structural ribosome constitution, respectively), may also contribute to an enhanced fitness of this strain. This may be particularly relevant considering that their mutational signature in the hybrid L2b/D-Da strain matches the one observed for prevalent clade T1 strains (i.e., strains with E/F-like genomic backbone) and that the CT677-CT680 region shows phylogenetic signals of tropism and prevalence (Ferreira et al. 2014) (Supplemental Figure S2C). For example, it was speculated that increased fitness at or around *ompA* (preventing recombination) may underlie the expansion of genotype E, which is the most succeeded worldwide genotype among *C. trachomatis* STIs (Nunes et al. 2009; Hadfield et al. 2017). Altogether, considering that: i) the hybrid outbreak strain displays a novel antigenic and adhesin fingerprint while retaining the genome-dispersed virulence signature of a classical epidemic LGV (L2b) lineage; ii) L2b strains are capable of dual tropism, i.e., infection of the rectal mucosa (leading to proctitis) and/or dissemination through mononuclear monocytes to inguinal nodes (leading to the bubonic disease LGV); and iii) the presumably parental genital strain belongs to T1 clade, which enrolls the most ecologically succeeded genital strains; one can straightforwardly hypothesize that the observed recombination-driven unique genomic backbone may have implications on the transmission, tissue tropism and pathogenic capabilities of the circulating hybrid outbreak-associated strain. However, more definitive information regarding the potential enhanced phenotypic skills of the hybrid strain must await detection and analysis of more isolates from around the world.

In summary, this study reports an ongoing outbreak involving a unique L2b variant (with potential modified fitness) in the context of the parental L2b worldwide epidemics. The risk sexual behavior (e.g., multiple partners and chemsex engaging) and potential international character of the involved sexual networks enhances the likelihood of this outbreak to be already spread abroad, which might be soon confirmed if broader *C. trachomatis* genotyping strategies are followed. The non-eligibility of this outbreak as laboratory-confirmed LGV cases additionally highlight the need to revise the criteria established by the European Commission, adopted by several countries, including Portugal, for LGV laboratory notification.

## Contributors

CF, AMR, JA, JA, JR, MR, RC were responsible for clinical diagnosis. DC, AIS, ZL, CC performed molecular diagnosis and *ompA* genotyping. LV supervised capillary and next-generation sequencing (NGS) procedures. JI did NGS procedures. VB did bioinformatics analysis. MJB, VB and JPG analysed and interpreted the data. VB and JPG wrote the Article. All authors revised and approved the Article. VB, MJB and JPG jointly supervised the study.

## Data access

*C. trachomatis-specific* reads (to avoid releasing reads matching to human genome) were deposited in the European Nucleotide Archive (ENA) (BioProject PRJEB32243). The assembled genome of a representative strain (Ct_L2b/D_PT05) of the hybrid L2b/D *C. trachomatis* outbreak strain was also deposited in ENA under the accession numbers ERZ870055 (sample ERS3372269 / SAMEA5570248).

## Declaration of interests

All other authors declare no competing interests.

---

**Supplemental Figure S1.**
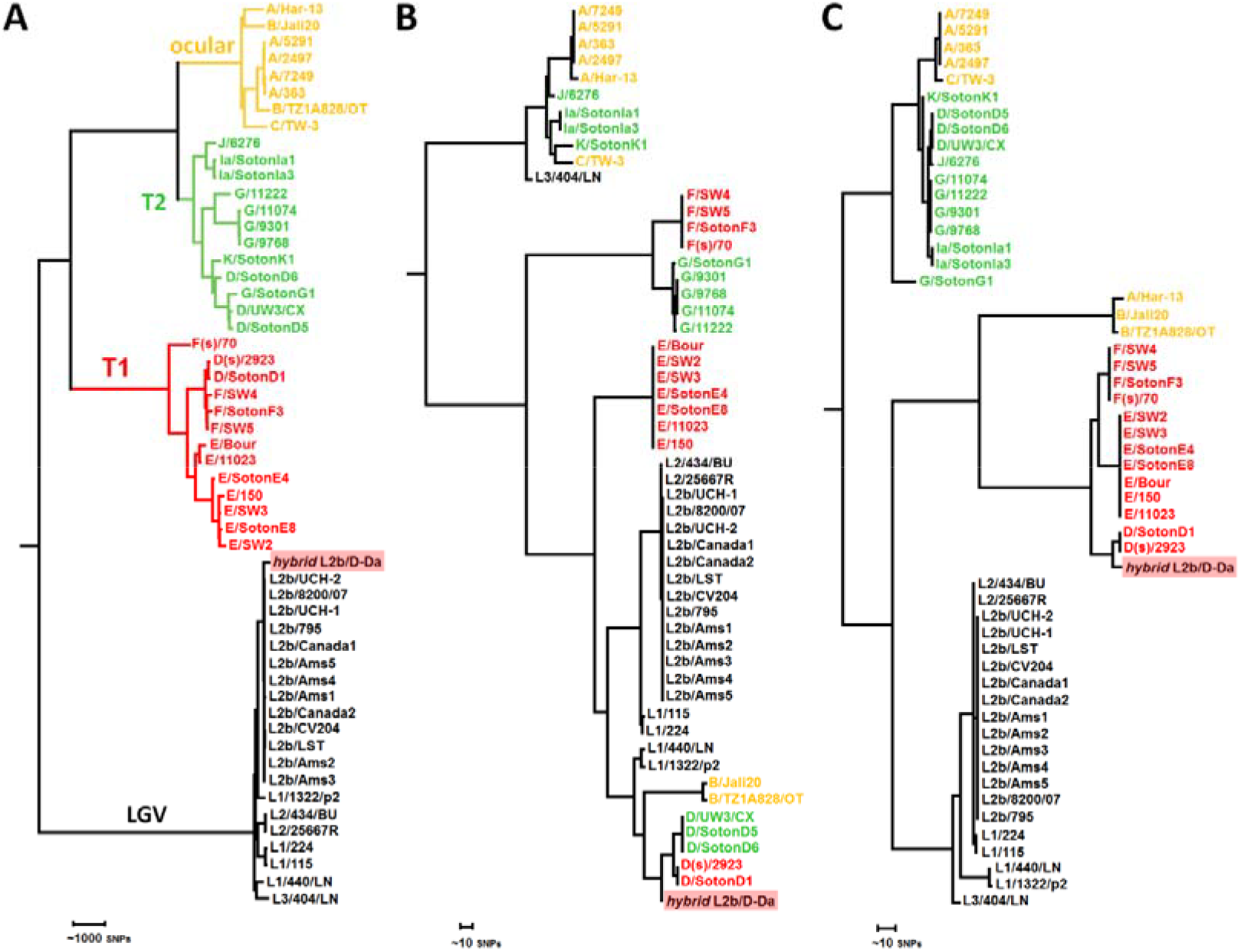
Phylogenetic integration of the hybrid L2b/D-Da outbreak-causing *C. trachomatis* strain in the genome-based species tree (A), *ompA* tree (B) and concatenated CT677-CT680-based tree (C). Represented strains reflect the species phylogenetic diversity (Harris et al. 2012) and are colored by the genome-based phylogenetic clade (ocular, prevalent genital - T1, non-prevalent genital - T2, and LGV) marked in panel A.

**Supplemental Figure S2.** *(see Excel file)*

**Supplemental Table S1.** *(see Excel file)*

**Supplemental Table S2.** *(see Excel file)*

